## Supplemental Figures and Tables for "Exceptional stability of a perilipin on lipid droplets depends on its polar residues, suggesting multimeric assembly"

**Figure S1**

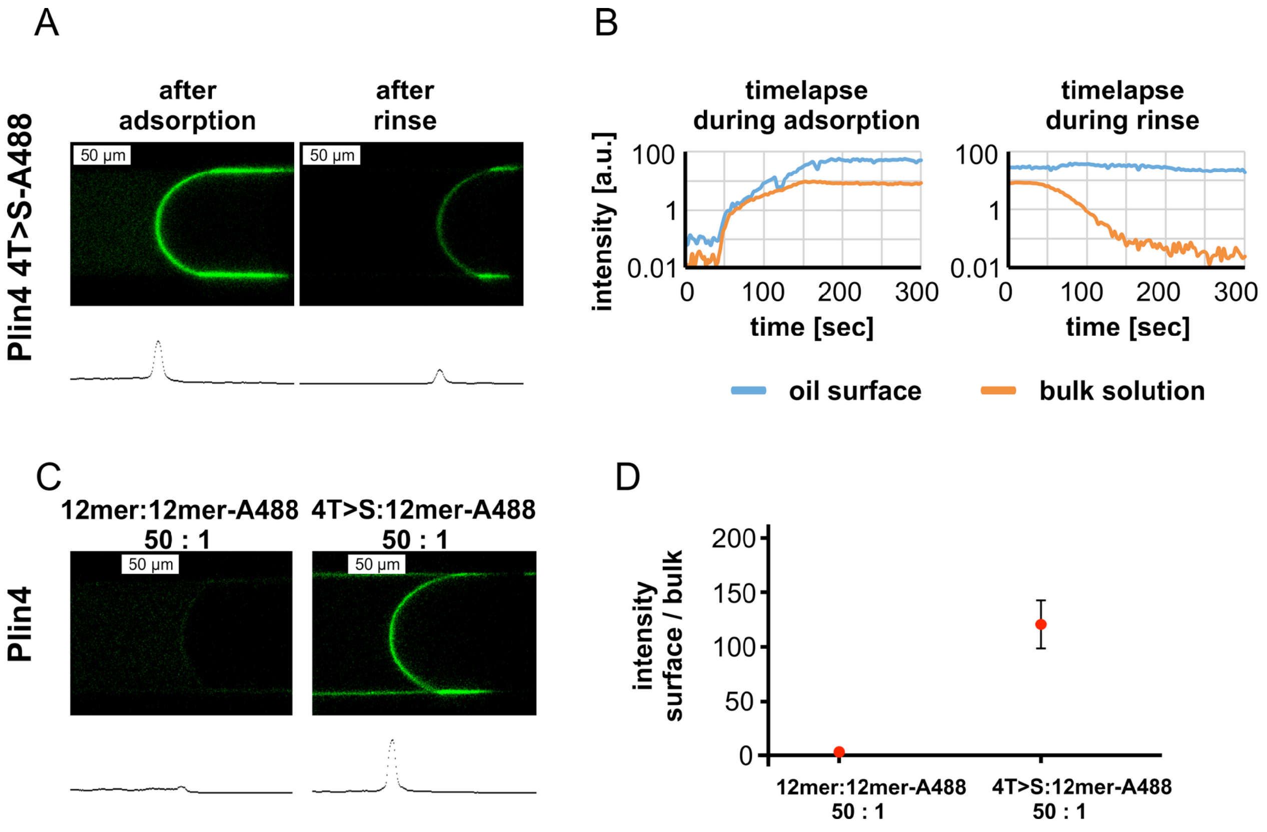


**Figure S1. Plin4 mutant 4T>S-A488 adsorbs irreversibly to the oil surface, but its interaction with oil is weaker than that of Plin4 12mer-A488. A.** Confocal images of the oil-buffer interface in the microfluidic system after adsorption of Plin4 4mer mutant 4T>S-A488 (0.1 mg/ml) on the oil surface and after rinsing with buffer. The intensity profile along the channel center is shown below each confocal image. **B.** Time evolution of signal intensity on the oil surface and in the bulk solution during adsorption and rinsing. **C.** Competition between Plin4 12mer and 4T>S mutant. The ratio between the intensity on the oil surface and in the bulk solution for a mixture of unlabeled Plin4 12mer : Plin4 12mer-A488 (50:1) and a mixture of unlabeled 4T>S : Plin4 12mer-A488 (50:1).

### Figure S2

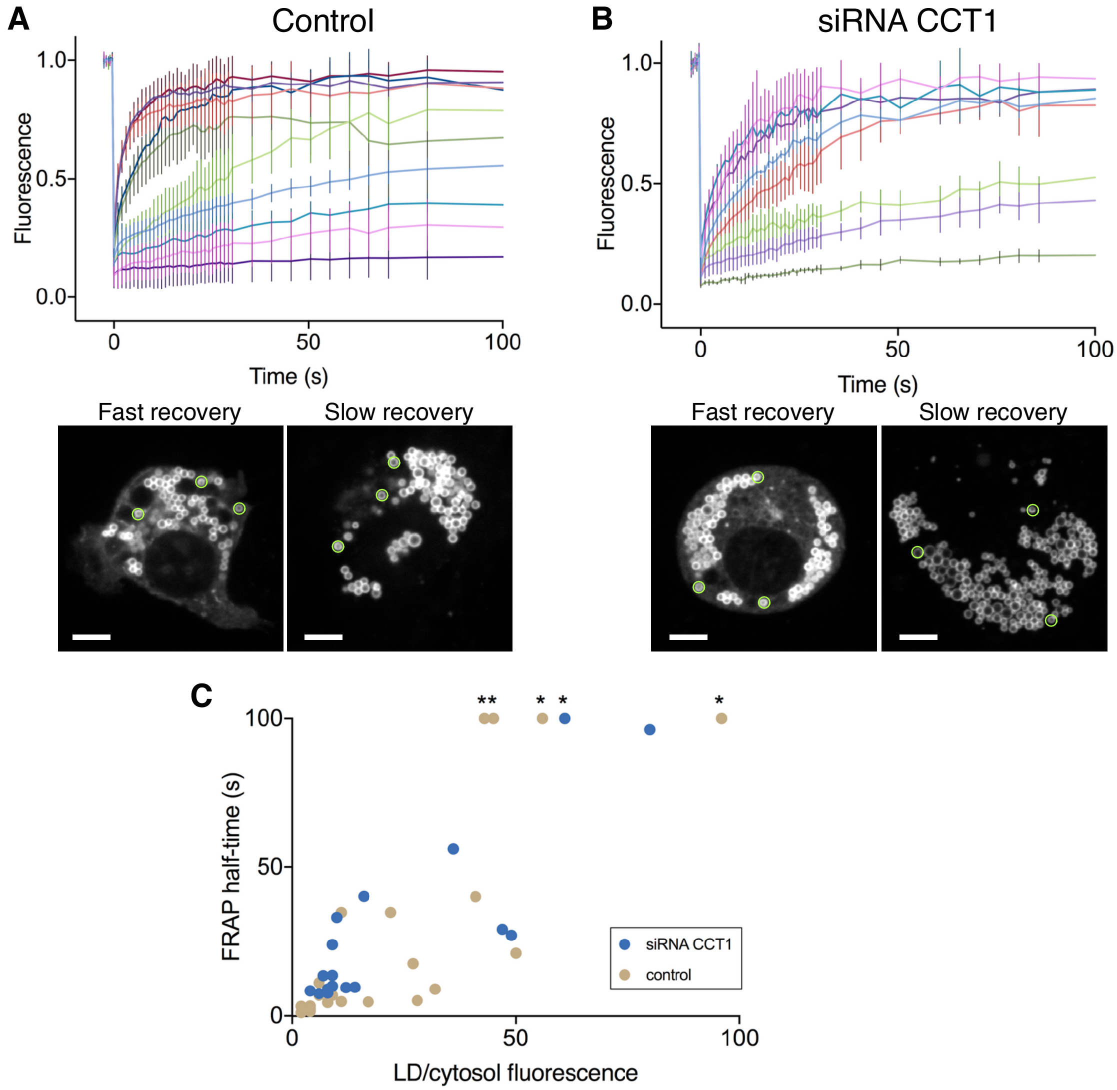


**Figure S2. Cell-to-cell variability in the recovery of Plin4 12mer-GFP after photobleaching of LDs in Drosophila S2 cells.** FRAP was performed on oleic acid-induced LDs in S2 cells stably transfected with Plin4 12mer-GFP. **A.** Graph shows fluorescence recovery curves from individual cells in the same experiment performed in control cells (RNAi against luciferase). Each curve represents mean ± SD from FRAP on 3 LDs in the same cell, as exemplified in the two images below the graph, showing two cells before bleaching. Areas that were bleached are marked with green circles. Fast fluorescence recovery was observed in the cell on the left, slow in the cell on the right. The signal intensity in the two images was adjusted to the same level, to show that the total Plin4-GFP expression level in the two cells was quite similar. However, note the difference in the amount of cytosolic signal. Scale bar: 5 m. **B.** Same as A, except that FRAP was performed on cells in which CCT1 has been depleted by RNAi. **C.** Graph showing correlation between the ratio of LD/cytosolic Plin4 12mer-GFP signal, and the half-time of recovery, determined from the curves shown in A and B. Each dot represents one cell, data is from two independent experiments. Asterisks denote cells which had very slow recovery kinetics (half-time > 100 s; in this case the value was manually set to 100 s.

**Figure S3**

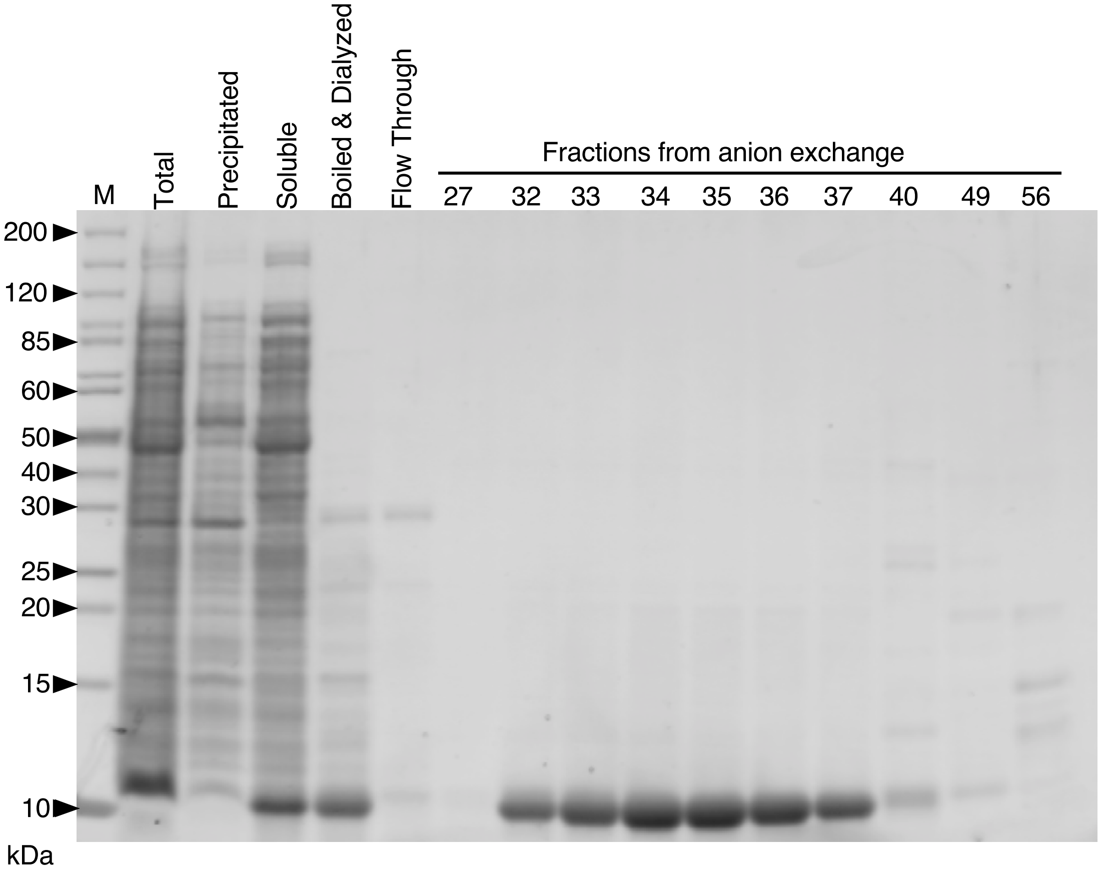


**Figure S3. Purification of Plin3 AH.** Tricine SDS-PAGE showing the purification of Plin3 AH after expression in *E. coli*. The indicated lanes show the proteins present in the total bacteria extract (total) and after a 4-step protocol, which includes centrifugation to separate the soluble fraction (Soluble) from the precipitated fraction (Precipitated), boiling and centrifugation of the soluble fraction to get the heat-resistant fraction, dialysis of the heat resistant fraction (boiled & dialyzed) and anion exchange chromatography with a gradient of NaCl from 10 to 1000 mM (numbered fractions). Gel stained with SyproOrange. Sizes of molecular weight protein standards are indicated.

**Figure S4**

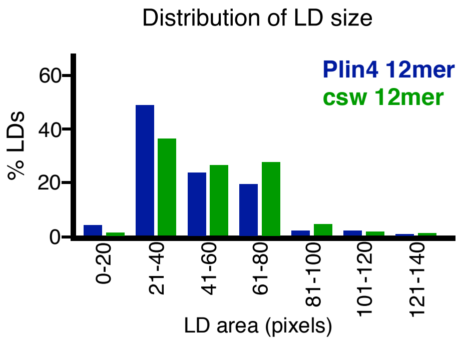


**Figure S4. Influence of redistribution of charge in Plin4 12mer on LD size in yeast.** Plot shows distribution of LD size in *pet10*Δ cells incubated with OA for 24h, and GFP fusions of Plin4 12mer or csw 12mer mutant. Graph shows a representative of three independent experiments, with 270 LDs measured for each construct. Pixel size: 0.091 um x 0.091 um.

**Supplementary Table 1. Analysis of Plin4 AH length in mammals.**

| Species | Uniprot entry code | AH length (aa) | # of repeats[1] | AH start (aa)[2] | AH end (aa)[2] | Repeat perturbations[3] | Plin4 length (aa) |
|---|---|---|---|---|---|---|---|
| *Mus musculus* | O88492 | 1022 | 31 | 105 | 1127 | Δ11 at 1075 | 1403 |
| *Ictidomys tridecemlineatus* | I3MUC2 | 819 | 25 | 107 | 926 | Δ12 at 310, +4 at 734 | 1291 |
| *Macaca mulatta* | F7FM35 | 899 | 27 | 20 | 919 | 0 | 1206 |
| *Homo sapiens* | Q96Q06 | 952 | 29 | 80 | 1032 | 0 | 1357 |
| *Callithrix jacchus* | F7GWD9 | 926 | 28 | 20 | 946 | 0 | 1200 |
| *Tupaia chinensis* | L8Y4I8 | 736 | 22 | 21 | 757 | Δ2 at 194 | 1210 |
| *Bos taurus* | F1MNM7 | 883 | 27 | 78 | 961 | Δ1 at 230 | 1260 |
| *Sus scrofa* | F1S7K4 | 978 | 30 | 107 | 1085 | Δ7 at 1080 | 1425 |
| *Ailuropoda melanoleuca* | G1L582 | 949 | 29 | 80 | 1029 | +2 at 560, +1 at 939 | 1343 |
| *Canis lupus familiaris* | J9JHP1 | 1169 | 35 | 104 | 1273 | 0 | 1588 |
| *Felis catus* | M3W5M0 | 971 | 29 | 82 | 1053 | +1 at 632 | 1336 |
| *Myotis davidii* | L5MGD7 | 1919 | 58 | 107 | 2026 | 0 | 2319 |
| *Myotis brandtii* | S7MI15 | 1544 | 47 | 115 | 1659 | 0 | 1957 |
| *Loxodonta africana* | G3UDZ0 | 1051 | 32 | 106 | 1157 | 0 | 1415 |
| *Monodelphis domestica* | F7DNE7 | 1096 | 33 | 20 | 1116 | 0 | 1339 |
| *Ornithorhynchus anatinus* | F7FVK1 | 842 | 26 | 14 | 946 | +90 at 590 | 1227 |

[1] The number of 33-aa repeats identified in each Plin4 sequence.

[2] The positions of the first and the last aa of the identified 33-aa repeats are indicated.

[3] (+) indicates insertion of aa (number indicated) between the 33-aa repeats at the indicated position in the Plin4 sequence. (Δ) indicates aa missing from a repeat. In all other cases, the 33-aa repeats are consecutive. (0) indicates that all repeats are consecutive.

**Supplementary Table 2. Plasmids used in this study.**

| Name | Insert | Region (aa) (1) | Vector | Host (2) | Source |
|---|---|---|---|---|---|
| pCLG03 | PLin4 4mer | hPlin4(246-377) | pET21b | E. coli | Copic 2018 |
| pKE23 | Plin4 12mer | hPlin4(510-905) | pET21b | E. coli | Copic 2018 |
| pSB49 | 4T>S (4mer) | 4x[246-278 M5t] (3) | pmCherry-N1 | Mamm | Copic 2018 |
| pMGA9 | 4T>S (4mer) | 4x[246-278 M5t] | pET21b | E. coli | This study |
| pGFP-Plin1 | Human Plin1 | Full cDNA | pGREG576 (ADH1pr, GFP) | Yeast | Jacquier 2013 |
| pGFP-Plin2 | Human Plin2 | Full cDNA | pGREG576 (ADH1pr, GFP) | Yeast | Jacquier 2013 |
| pGFP-Plin3 | Human Plin3 | Full cDNA | pGREG576 (ADH1pr, GFP) | Yeast | Jacquier 2013 |
| pRHT140 | ADHpr-mcs-GFP | | pRS416 (CEN-URA3) | Yeast | S. Leon |
| pMGA4 | ADHpr-mcs-mCherry (swap of GFP in pRHT140) | | pRS416 (CEN-URA3) | Yeast | This study |
| pMGA10 | Plin1 AH-GFP | hPlin1(aa108-194) | pRHT140 | Yeast | This study |
| pMGA5 | Plin2 AH-GFP | hPlin2(aa100-192) | pRHT140 | Yeast | This study |
| pMGA7 | Plin3 AH-GFP | hPlin3(aa113-205) | pRHT140 | Yeast | This study |
| pMGA28 | Plin3(87-205)-GFP | hPlin3(aa87-205) | pRHT140 | Yeast | This study |
| pMGA19 | Plin3 AH | hPlin3(aa113 – 205) | pET21b | E. coli | This study |
| pKE31 | Plin4 4mer-GFP | hPlin4(aa246-377) | pRHT140 | Yeast | Copic 2018 |
| pKE33 | Plin4 12mer-GFP | hPlin4(aa510-905) | pRHT140 | Yeast | Copic 2018 |
| pMGA16 | Plin4 12mer-mCherry | hPlin4(aa510-905) | pMGA4 | Yeast | This study |
| pCLG26 | Plin4 8mer | hPlin4(aa246-509) | pmCherry-N1 | Mamm | Copic 2018 |
| pMGA30 | Plin4 6mer-GFP | hPlin4(aa246-433) | pRHT140 | Yeast | This study |
| pMGA22 | Plin4 8mer-GFP | hPlin4(aa246-509) | pRHT140 | Yeast | This study |
| pACJ22 | Plin4 4mer | hPlin4(aa246-377) | pmCherry-N1 | Mamm | Copic 2018 |
| pSB58 | 2D>E (4mer) | 4x[246-278 M17e] | pmCherry-N1 | Mamm | This study |
| pSB60 | NN (4mer) | 4x[246-278 M18q] | pmCherry-N1 | Mamm | This study |
| pSB83 | QN (4mer) | 4x[246-278 M19qn] | pmCherry-N1 | Mamm | This study |
| pSB65 | QQ (4mer) | 4x[246-278 M20q2] | pmCherry-N1 | Mamm | This study |
| pSB86 | 3K>R | 4x[246-278 M21r] | pmCherry-N1 | Mamm | This study |
| pCLG62 | Plin4 12mer | hPlin4(aa510-905) | pmCherry-N1 | Mamm | Copic 2018 |
| pSB06 | Csw 12mer | Charge swap of hPlin4(aa510-905) (3) | pmCherry-N1 | Mamm | This study |
| pSB41 | Plin4 12mer-GFP | hPlin4(aa510-905) | pMTWG | Dros. | Copic 2018 |
| pMGA1 | Csw 12mer | Charge swap of hPlin4(aa510-905) | pET21b | E. coli | This study |

(1) Position of amino acids (aa) in human perilipin sequences..
